# Establishing the fluorescence-activating and absorption-shifting tag as a fluorescent reporter protein in *Methanothermobacter thermautotrophicus* ΔH

**DOI:** 10.64898/2026.05.29.728764

**Authors:** Tina Baur, Maximilian Flaiz, Largus T. Angenent, Bastian Molitor

**Affiliations:** Environmental Biotechnology Group, Department of Geosciences, University of Tübingen, Tübingen, Germany; Microbial Metabolic Biochemistry, Institute of Biochemistry, University of Leipzig, Leipzig, Germany; Cluster of Excellence – Controlling Microbes to Fight Infections, University of Tübingen, Tübingen, Germany; Max Planck Institute for Developmental Biology, Tübingen, Germany; Department of Biological and Chemical Engineering, Aarhus University, Aarhus, Denmark; The Novo Nordisk Foundation CO_2_ Research Center (CORC), Aarhus University, Gustav Wieds Vej 10C, 8000 Aarhus C, Denmark

**Keywords:** *Methanobacterium thermoautotrophicum*, promoter activities, fluorescence-activating and absorption-shifting tag, reporter protein, anaerobic thermophile, methanogen, archaea

## Abstract

The thermophilic methanogen *Methanothermobacter thermautotrophicus* ΔH is a model microbe for hydrogenotrophic methanogenesis and an emerging platform host for metabolic engineering. Despite recent advances in its genetic accessibility, the available molecular toolbox lacks fluorescent reporter proteins that are suitable for anaerobic and thermophilic conditions. Here, we established the fluorescence-activating and absorption-shifting tag (FAST) as a reporter protein in *M. thermautotrophicus*. We expressed codon-optimized variants of FAST by applying established genetic tools, and evaluated the performance for two temperatures and three fluorogens. We demonstrated that FAST is functional in *M. thermautotrophicus* but exhibits temperature-dependent instability, which is more pronounced at 60°C compared to 50°C. Among the tested fluorogens, ^TF^Lime and ^TF^Amber yielded comparable fluorescence intensities, while ^TF^Coral resulted in significantly lower fluorescence intensity. Exploiting the partial thermolability of FAST, we characterized the dynamic expression profiles of several promoters, which revealed growth phase-dependent regulation patterns. Our findings challenge previous assumptions of constitutive expression for several promoters. Notably, we identified distinct expression patterns for promoters that are associated with methanogenesis and energy-converting hydrogenases. Our results establish FAST as a versatile fluorescent reporter for thermophilic methanogens and provide new insights into promoter regulation in *M. thermautotrophicus*. This work expands the genetic toolbox for this microbe and lays the foundation for advanced studies in archaeal cell biology and biotechnology.

## 1 Introduction

The thermophilic methanogenic archaeon (methanogen) *Methanothermobacter thermautotrophicus* ΔH (hereafter *M. thermautotrophicus*) is a well-established model microbe for studying hydrogenotrophic methanogenesis (Costa and Whitman, 2023). Despite decades of intensive research, this strain has only recently become genetically tractable (Fink et al., 2021). Its genetic toolbox comprises replicating shuttle plasmids, constitutive and inducible promoter systems, and genome engineering tools that are based on allelic exchange (Fink et al., 2023; Baur et al., 2026). Just recently the recombinant production of acetoin has been demonstrated, highlighting its potential for biotechnological applications beyond methane production (Pfeifer et al., 2021; Contreras et al., 2022; Baur et al., 2026). Although these recent advances enable many applications, the molecular toolbox of *M. thermautotrophicus* lacks fluorescent reporter proteins, which are powerful tools for studying gene expression, protein localization, and protein-protein interactions (Thorn, 2017; Flaiz and Sousa, 2024).

The discovery of the green fluorescent protein (GFP) from the jellyfish *Aequorea victoria* and its recombinant production has revolutionized many disciplines, including molecular genetics and cell biology, and was awarded the Nobel prize in Chemistry in 2008 (Shimomura et al., 1962; Chalfie et al., 1994). Today fluorescent reporter proteins are an indispensable tool in molecular biology and have transformed the capacities in studying cellular processes in all domains of life (Specht et al., 2017; Bisson-Filho et al., 2018; Charubin et al., 2018). However, as the maturation of the intrinsic fluorophore of GFP and its many derivatives is oxygen-dependent, their use is limited in anaerobic microbes, such as *M. thermautotrophicus* (Charubin et al., 2018). Over the past years, several alternative fluorescent reporters have been developed for use in anoxic environments. For example, ligand-independent flavin-based fluorescent proteins (FbFPs) (Drepper et al., 2007) have been successfully applied as reporters in various strictly anaerobic *Clostridia* (Streett et al., 2021). Although their ligand independence is a major advantage, their application is limited by relatively low fluorescence intensity and, more importantly, their emission is restricted to the cyan-green spectrum. This is particularly problematic for use in methanogens, such as *M. thermautotrophicus*, because the cofactor F_420_ exhibits strong intrinsic autofluorescence in the same spectral range (Lambrecht et al., 2017). This limitation can be overcome by harnessing ligand-dependent fluorescent proteins, including the HaloTag, SNAP-tag, and the fluorescence-activating and absorption-shifting tag (FAST). These proteins depend on fluorogenic ligands (fluorogens) that can be tuned to different spectral properties, enabling the bypass of cellular autofluorescence. Both HaloTag and SNAP-tag show strong fluorescence in the anaerobic bacteria *Clostridium acetobutylicum* and *Clostridium ljungdahlii*, but their externally supplied ligand binds covalently to the protein and requires long labeling times, limiting their use for real-time studies (Charubin et al., 2020). As both the HaloTag and SNAP-tag originate from mesophilic sources (Keppler et al., 2003; Los et al., 2008) their functionality under thermophilic conditions remains uncertain.

FAST provides a ligand-dependent alternative that relies on external fluorogens that do not bind covalently. The binding of the fluorogens is specific, rapid, fully reversible, and can, thus, be dynamically exchanged (Gautier, 2022). Among the different reporters, FAST resulted in fluorescence in several acetogenic and solventogenic bacteria, including the mesophilic species *C. ljungdahlii, Eubacterium limosum, Acetobacterium woodii, C. acetobutylicum*, and *Clostridium saccharoperbutylacetonicum* (Streett et al., 2019; Charubin et al., 2020; Baur et al., 2022; Mook et al., 2022; Flaiz et al., 2024). FAST was engineered from the mesophilic photoactive yellow protein of *Halorhodospira halophila* (Plamont et al., 2016). However, an evolved variant of FAST, promiscuous FAST (pFAST), showed superior chromophore binding and fluorescence brightness properties compared to the original FAST (Benaissa et al., 2021), and further adaptive laboratory evolution enabled its use in the thermophilic species *Thermoanaerobacter kivui* (Hocq et al., 2023; Sitara et al., 2025). Beyond the domain Bacteria, FAST has also been successfully implemented in the domain Archaea, specifically in the mesophilic methanogens *Methanococcus maripaludis* and *Methanosarcina acetivorans*, showing its functionality on a single-cell level for live-cell imaging and flow cytometry, respectively (Hernandez and Costa, 2022; Adlung and Scheller, 2023). Therefore, FAST represents a promising fluorescent reporter protein for the application as a molecular tool in thermophilic methanogens.

Here, we report the implementation of FAST in the thermophilic methanogen, *M. thermautotrophicus*, adding to the list of anaerobic and thermophilic microbes for which FAST has been established. We applied our previously established genetic system (Fink et al., 2021) to implement FAST as a fluorescent reporter protein into the available toolbox. This enabled us to identify a limited temperature stability in our system, evaluate the functionality of different fluorogens, and eventually study the dynamic expression profile of several promoters.

## 2 Materials and Methods

### 2.1 Microbial strains and cultivation

Microbial strains and plasmids that were used in this study are listed in **Table 1**. *Escherichia coli* NEB stable was used for cloning purposes, and *E. coli* S17-1 was used for interdomain conjugational DNA transfer into *M. thermautotrophicus* (Fink et al., 2021). *E. coli* strains were cultivated aerobically in lysogeny broth (LB) medium (Green et al., 2012) at 37°C with proper antibiotic supplementation and shaking (150 rpm). Cells of *M. thermautotrophicus* were grown anaerobically in liquid or on solid mineral salt (MS) medium as previously described (Fink et al., 2021). Antibiotics were supplemented for the selection of recombinant strains to the following concentrations: 30 µg mL^−1^ chloramphenicol; 250 µg mL^−1^ neomycin; 100 µg mL^−1^ carbenicillin; and 10 µg mL^−1^ trimethoprim. Growth experiments with recombinant *M. thermautotrophicus* strains were conducted in 50 mL MS medium with neomycin in 250-mL serum bottles. In all cases, the headspace of the serum bottles was filled with an H_2_/CO_2_ (80:20 vol-%) mixture to 1 bar overpressure. Cultures were grown at 50°C or 60°C under constant shaking (150 rpm), and growth was monitored by measurement of the optical density at 600 nm (OD_600_; BioMate™ 160 UV-vis spectrophotometer, Thermo Fisher Scientific, Waltham, MA, USA). To determine the fluorescence of FAST-producing cells, 4-8 mL of culture was withdrawn at different times of growth and processed as described in section 2.3.

**Table 1.**
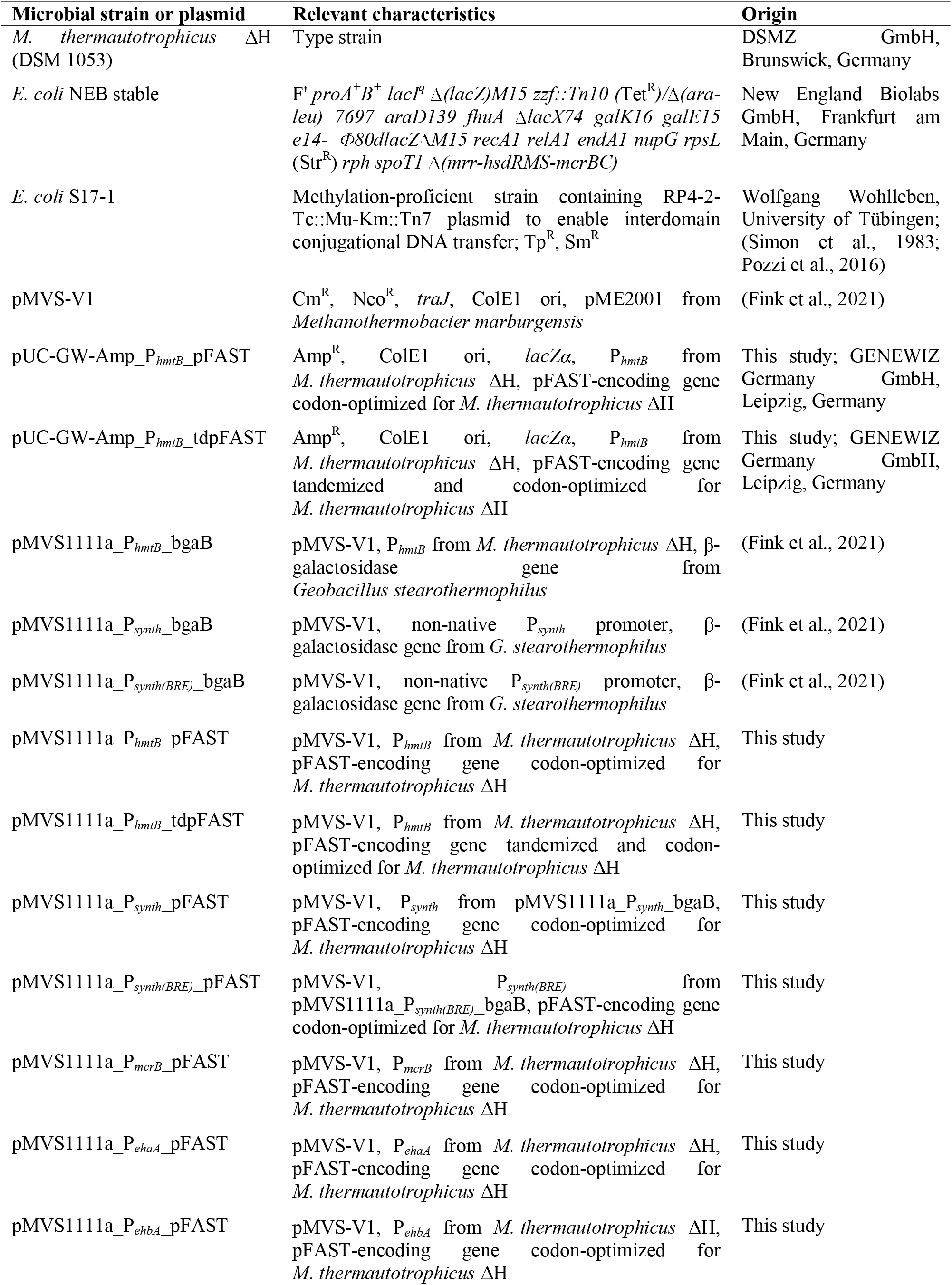
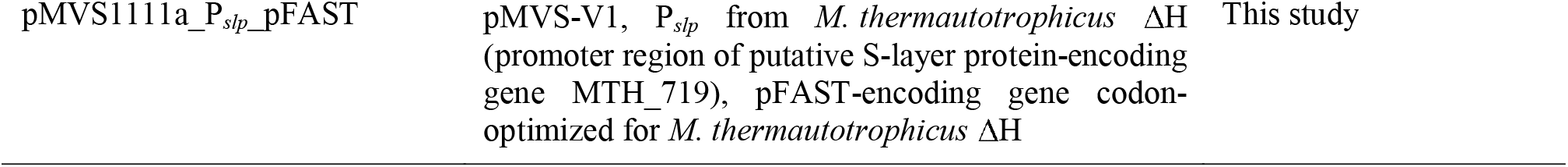
Microbial strains and plasmids used in this study.

### 2.2 Plasmid construction and DNA transfer

Standard molecular techniques were applied for plasmid construction. All primers (Integrated DNA Technologies, Coralville, IA, USA) that were used in this study are summarized in **Table 2**. DNA fragments for cloning purposes were generated *via* PCR using the Q5 High-Fidelity DNA Polymerase (New England Biolabs, Ipswich, MA, USA) or *via* restriction digestion with restriction enzymes from New England Biolabs (Ipswich, MA, USA). Generated DNA fragments were purified from agarose gels with the Zymoclean™ Gel DNA Recovery Kit (ZYMO Research Corp., Irvine, CA, USA), according to the manufacturer’s instructions. All plasmids were constructed *via* Gibson assembly using the NEBuilder® HiFi DNA Assembly Master Mix or *via* classic restriction/ligation using the T4 DNA Ligase (New England Biolabs, Ipswich, MA, USA) with matching restriction sites. Subsequently, chemically competent *E. coli* NEB stable cells were transformed with ligation/assembly mixtures as previously described (Fink et al., 2021). Successful plasmid constructions were verified *via* restriction digestion and Sanger sequencing (GENEWIZ Germany GmbH, Leipzig, Germany) of prepared plasmid DNA from individual *E. coli* NEB stable colonies (Zyppy™ Plasmid Miniprep Kit, ZYMO Research Corp., Irvine, CA, USA).

**Table 2.**
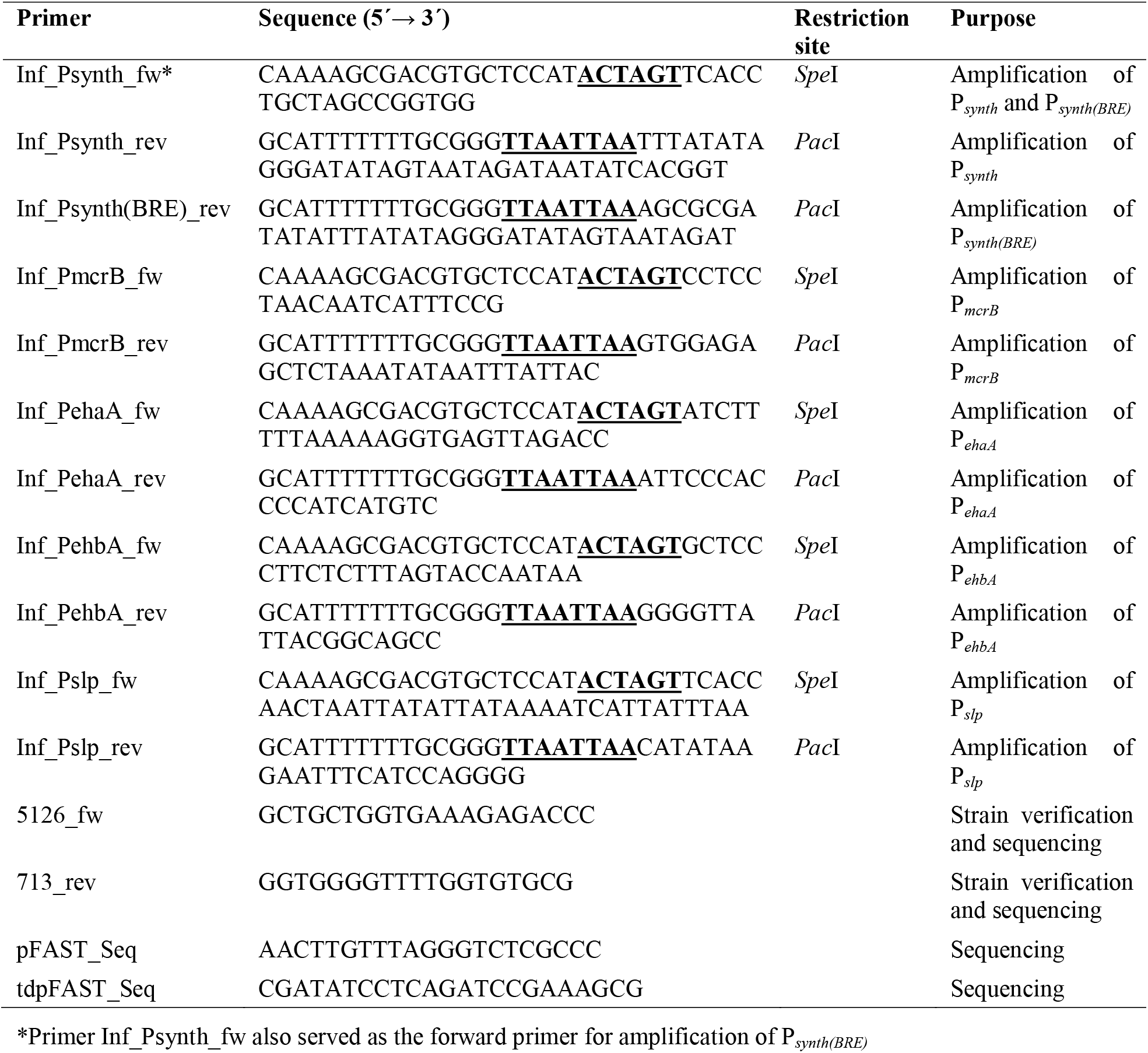
Primers used in this study. Restriction sites are underlined in bold.

For the construction of plasmids, pFAST (Benaissa et al., 2021) and a tandemized version, which is called tandem-pFAST (tdpFAST; two pFAST connected by a glycine linker [GGGSGGG]), were codon-optimized for *M. thermautotrophicus* applying the GENEius light software tool (Eurofins Genomics Europe Shared Services GmbH, Ebersberg, Germany; **Supplementary Table 1**). Acquired gene sequences were placed under the control of the P_*hmtB*_ promoter from *M. thermautotrophicus* (Fink et al., 2021). Synthesis, codon-optimization, and subcloning of *in silico*-generated P_*hmtB*_-pFAST and P_*hmtB*_-tdpFAST fragments in pUC vectors were performed by GENEWIZ Germany GmbH (Leipzig, Germany), yielding pUC-GW-Amp_P_*hmtB*__pFAST and pUC-GW-Amp_P_*hmtB*__tdpFAST. To construct the initial FAST plasmids, pMVS1111a_P_*hmtB*__bgaB was linearized using *Asc*I and *Pac*I. The eliminated *bgaB* gene was replaced with P_*hmtB*_-pFAST and P_*hmtB*_-tdpFAST inserts, which were cut out of pUC vectors using the same enzymes, resulting in plasmids pMVS1111a_P_*hmtB*__pFAST and pMVS1111a_P_*hmtB*__tdpFAST, respectively. pMVS1111a_P_*hmtB*__pFAST then served as the backbone for the assembly of all other plasmid variants *via* exchange of the promoter module, which was flanked by introduced *Pac*I and *Spe*I restriction sites. For this purpose, pMVS1111a_P_*hmtB*__pFAST was linearized using the respective enzymes and used as the backbone for all further cloning steps. Promoters to be characterized (**Supplementary Table 1**) were amplified using designed primers (**Table 2**) and pMVS1111a_P_*synth*__bgaB (P_*synth*_), pMVS1111a_P_*synth(BRE)*__bgaB (P_*synth(BRE)*_), or genomic DNA of *M. thermautotrophicus* ΔH (P_*mcrB*_, P_*ehaA*_, P_*ehbA*_, P_*slp*_) as the template DNA. A schematic plasmid map of the basic pFAST vector is provided in **Supplementary Figure 1**.

Constructed plasmids were first transferred into *E. coli* S17-1 and subsequently into *M. thermautotrophicus*, according to the procedures described by Fink et al. 2021 with minor modifications (Fink et al., 2021). Instead of step-wise centrifugation of 8 mL *M. thermautotrophicus* culture in the anaerobic chamber, 20 mL of early-stationary culture was transferred into a 50-mL Falcon tube and then centrifuged outside the chamber at room temperature for 10 min at 3,684 x g. Afterwards, cells were transferred back into the anaerobic chamber and resuspended in 600 µL of the supernatant medium. Subsequently, 300 µL of concentrated *M. thermautotrophicus* suspension was gently mixed with the pellet of 10 mL of the respective *E. coli* S17-1 carrying the plasmid to be conjugated. Finally, 100 µL of the *M. thermautotrophicus*-*E. coli* S17-1 cell suspension was spotted on LB-MS agar for mating as previously described (Fink et al., 2021), with the difference that (per Liter) 10 g peptone and 5 g yeast extract were added to the MS medium during preparation. Recombinant strains were verified *via* PCR (Phire Plant Direct PCR Master Mix, Thermo Fisher Scientific, Waltham, MA, USA) after picking single colonies from MS agar plates and growing them in liquid cultures (Fink et al., 2021) with primers 5126_fw and 713_rev (**Table 2**). Amplified DNA fragments were sent for Sanger sequencing (GENEWIZ Germany GmbH, Leipzig, Germany) to confirm mutation-free recombinant *M. thermautotrophicus* strains.

### 2.3 FAST assays

To evaluate the activities of different promoters in *M. thermautotrophicus*, FAST-mediated fluorescence was monitored throughout growth. For that, 4-8 mL of culture was withdrawn at different stages of growth and processed as previously described with minor adjustments (Flaiz et al., 2021). Cells were harvested aerobically (10 min, 3,684 x g, 4°C) and then washed once with cold phosphate-buffered saline (PBS, 137 mM NaCl, 10 mM Na_2_HPO_4_ x 2 H_2_O, 2 mM KH_2_PO_4_, 2.7 mM KCl, pH 7.4). After another centrifugation step (10 min, 3,684 x g, 4°C), cells were suspended in PBS to a final OD_600_ of 1. Fluorescence intensities (FLU) of the cultures were determined using a microplate reader (Infinite 200 pro M Nano+, Tecan Deutschland GmbH, Crailsheim, Germany). For this, 100 µL of cell suspension was transferred to black, flat-bottomed 96-well plates (Greiner Bio-One GmbH, Frickenhausen, Germany) and supplemented with one of the fluorescent dyes ^TF^Lime, ^TF^Amber, or ^TF^Coral (The Twinkle Factory, Paris, France) to a final concentration of 10 µM. Cells without the addition of the fluorogens served as the negative control for autofluorescence; respective values were subtracted from FLU values with supplementation of the fluorogens. Fluorescence was determined using the respective excitation and emission maxima of the FAST-fluorogen complex (^TF^Lime, ex. 480 nm/em. 541 nm; ^TF^Amber, ex. 499 nm/em. 558 nm; ^TF^Coral, ex. 516 nm/em. 600 nm). Finally, FLU values were normalized to the OD_600_ of PBS-suspended cells.

### 2.4 Statistical analysis

For conducting statistical tests, the samples were first analyzed for equal distribution with Shapiro-Wilk’s test. Because of the sample size of N = 3, a test for equal variance was omitted, and instead a more robust test towards potentially heterogenous variances between data sets was selected *a priori*. For the differences between empty-vector negative control, pFAST, and tdpFAST at 50°C and 60°C, respectively, a Welch’s analysis of variance (ANOVA) with Games-Howell *post-hoc* test was performed. For the comparison between the two temperatures for the two different groups, pFAST and tdpFAST, a Welch’s t-test with Holm correction was used. To compare the difference of the three fluorophores with pFAST, a one-factorial ANOVA with Tukey honest significant difference (HSD) *post-hoc* test was performed. For the comparison of the maximum fluorescence intensities between the different promoters, again a Welch’s ANOVA with Games-Howell *post-hoc* test was used.

## 3 Results

### 3.1 FAST can be used as a fluorescent reporter in *M. thermautotrophicus*

We intended to establish a fluorescent reporter protein in the thermophile *M. thermautotrophicus* to expand the capacities of the molecular toolbox for this microbe. We had previously established a thermostable β-galactosidase, which provides a valuable tool for molecular and synthetic biology (Fink et al., 2021; Baur et al., 2026). However, fluorescent reporter proteins have distinct benefits for the use in cell biology, especially, when fluorescence is enabled *in vivo* under thermophilic and anoxic conditions. Therefore, we sought to establish the FAST system, which recently was shown to be functional in the thermophilic bacterium *T. kivui* (Hocq et al., 2023). Hocq et al., 2023, described a certain thermolability in their system. Thus, we initially compared the two FAST variants, pFAST and tdpFAST, at temperatures of 50°C and 60°C. We used codon-optimized pFAST and tdpFAST versions for *M. thermautotrophicus* and gene expression was under the control of the P_*hmtB*_ promoter. We included *M. thermautotrophicus* [pMVS-V1] as the empty-vector negative control (**Figure 1, Supplementary Table 2**). We selected ^TF^Lime for this initial experiment, because this fluorogen has a GFP-like emission and excitation range. ^TF^Lime was also successfully applied in *T. kivui*, and the mesophilic methanogens *M. maripaludis* und *M. acetivorans*, sufficiently overcoming autofluorescence of the cells (Hernandez and Costa, 2022; Adlung and Scheller, 2023; Hocq et al., 2023). For all strains, growth was comparable and no growth differences were observed in the experiment (**Figure 1A**). For both variants, the maximum fluorescence intensity was significantly higher at 50°C compared to 60°C (pFAST, p = 0.002; tdpFAST, p = 0.0001; **Supplementary Table 2**). Furthermore, the fluorescence intensity was not stable but constantly decreased over time. This effect was less pronounced at 50°C, where residual fluorescence remained even after 48 h (**Figure 1B**). However, at 60°C, no fluorescence was detectable anymore after 24 h (**Figure 1C**). We found that pFAST led to a significantly higher maximum fluorescence intensity compared to tdpFAST at both temperatures (50°C, p = 0.027; 60°C, p = 0.001; **Figure 1D, Supplementary Table 2**). Thus, we confirmed the thermolability that was described for *T. kivui* (Hocq et al., 2023), and found this to be slightly more pronounced in our system. Based on these results, we performed all following experiments with an incubation temperature of 50°C.

**Figure 1.**
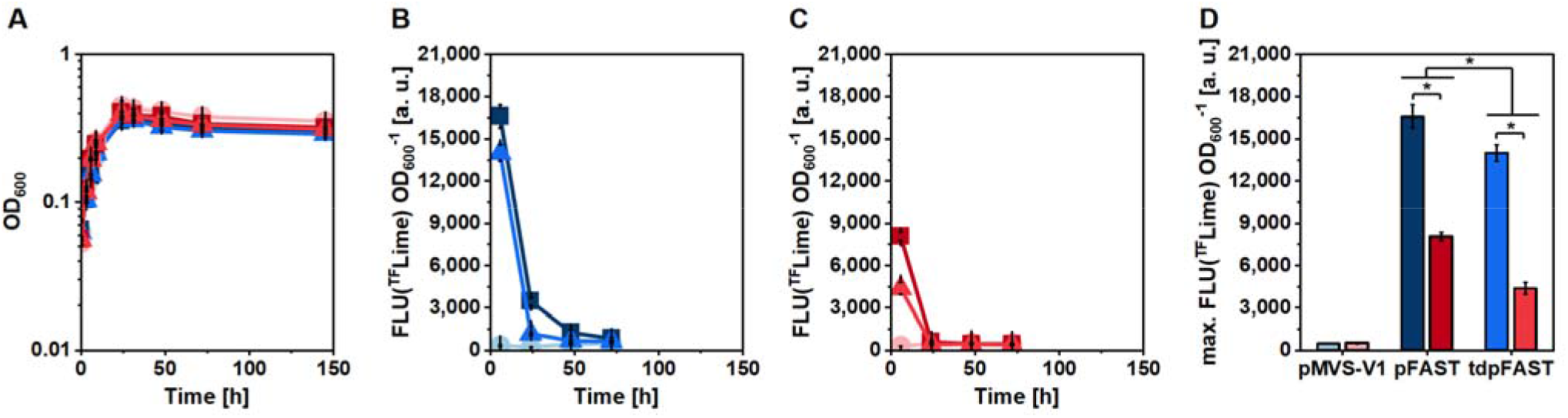
Establishment of FAST in *M. thermautotrophicus* ΔH. **(A)**, Growth (OD_600_); **(B)**, normalized fluorescence intensity (FLU(^TF^Lime) OD_600_ ^−1^) for cultivation at 50°C; **(C)** FLU(^TF^Lime) OD_600_ ^−1^ for cultivation at 60°C; **(D)**, maximum FLU(^TF^Lime) OD_600_ ^−1^ of FAST-producing *M. thermautotrophicus* strains. *, significant difference (p ≤ 0.05). Circles, *M. thermautotrophicus* [pMVS-V1]; squares, *M. thermautotrophicus* [pMVS1111a_P_*hmtB*__pFAST]; triangles, *M. thermautotrophicus* [pMVS1111a_P_*hmtB*__tdpFAST]. Blue, cultivation at 50°C; red, cultivation at 60°C. Error bars indicate standard deviations, N=3.

### 3.2 Different fluorogens can be applied with FAST in *M. thermautotrophicus*

We hypothesized that due to the autofluorescence of cofactor F_420_ (ex. 420 nm/em. 470-480 nm) (Lambrecht et al., 2017), higher fluorescence signals might be reached with the fluorogens ^TF^Amber or ^TF^Coral, because the excitation and emission wavelengths would interfere less with the autofluorescence signal. Therefore, we compared the performance of ^TF^Lime, ^TF^Amber, and ^TF^Coral with pFAST, under the control of the P_*hmtB*_ promoter at 50°C (**Figures 2 and 3**). Again, we did not observe an impact on the growth of the different strains (**Figure 2A**). ^TF^Amber resulted in slightly higher fluorescence intensities (**Figure 2C**) and less background (**Figure 3B**) than ^TF^Lime (but not significant, p = 0.573; **Figures 2B** and **3A**), while ^TF^Coral resulted in significantly lower fluorescence intensities (*vs*. ^TF^Lime, p = 0.009; *vs*. ^TF^Amber, p = 0.003; **Figures 2D** and **3C, Supplementary Table 2**). In conclusion, ^TF^Lime and ^TF^Amber gave comparable results and will be similarly effective in future work, allowing to select the most appropriate excitation/emission range, depending on the application.

**Figure 2.**
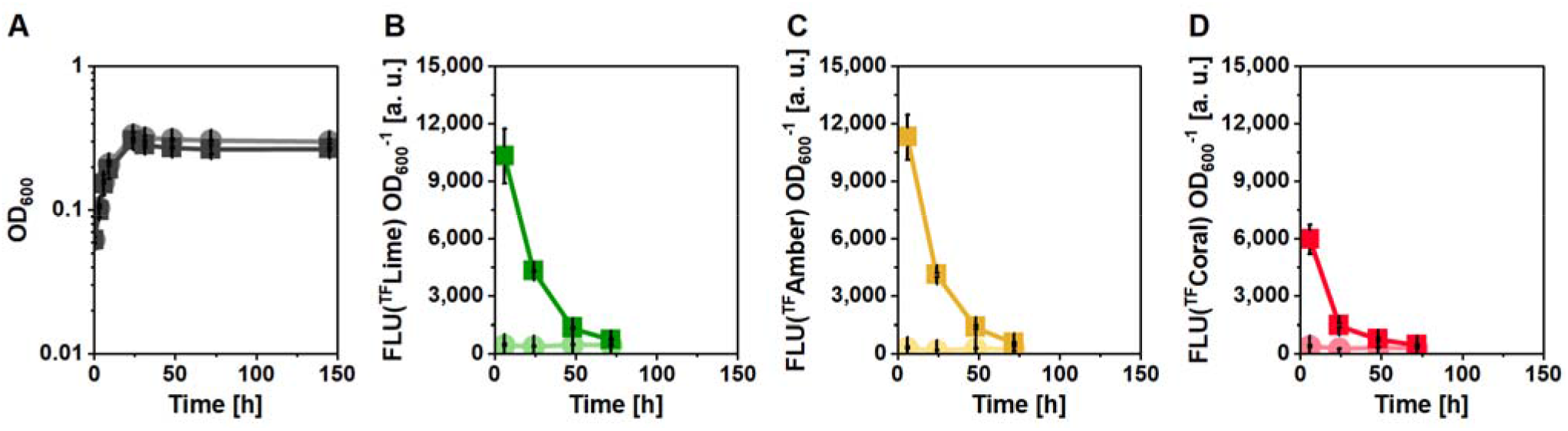
Characterization of different fluorogens in *M. thermautotrophicus* ΔH with pFAST at 50°C. **(A)**, Growth (OD_600_); **(B)**, normalized fluorescence intensity (FLU(^TF^Lime) OD_600_ ^−1^); **(C)**, FLU(^TF^Amber) OD_600_ ^−1^; **(D)**, FLU(^TF^Coral) OD_600_ ^−1^. Circles, *M. thermautotrophicus* [pMVS-V1]; squares, *M. thermautotrophicus* [pMVS1111a_P_*hmtB*__pFAST]. Error bars indicate standard deviations, N=3.

**Figure 3.**
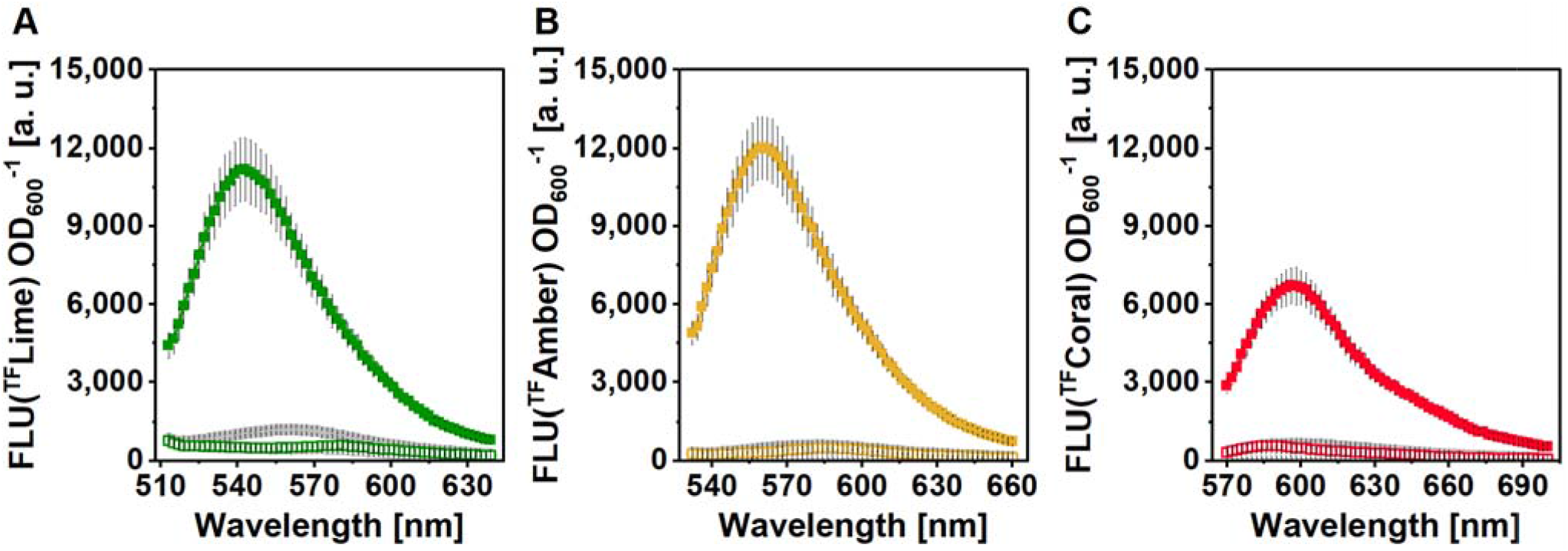
Emission spectra of pFAST-producing cells of *M. thermautotrophicus* ΔH after 24 h at 50°C. **(A)**, ^TF^Lime; **(B)**, ^TF^Amber; **(C)**, ^TF^Coral. Grey, *M. thermautotrophicus* [pMVS-V1]; green/yellow/red, *M. thermautotrophicus* [pMVS1111a_P_*hmtB*__pFAST]; open symbols, fluorescence intensities without fluorogen; filled symbols, fluorescence intensities with fluorogen. Error bars indicate standard deviations, N=3.

### 3.3 FAST allows the characterization of expression profiles of different promoters

We had previously considered the P_*hmtB*_ promoter to be constitutive, which was based on measurements with the thermostable β-galactosidase (Fink et al., 2021). However, our findings here indicated towards a more regulated expression profile. Therefore, we decided to test a small promoter library, including promoters that we had previously investigated (Fink et al., 2021) and an additional selection (**Table 3, Figure 4**), to understand their expression profiles in more detail. To increase comparability, we performed one experiment with all promoters, including pMVS-V1 as an empty-vector negative control, at 50°C with ^TF^Amber as the fluorogen (**Figure 5, Supplementary Table 2**).

**Table 3.** Overview of promoters characterized in this study using pFAST.

| Promoter | Origin (Locus tag) | Features | Reference |
| --- | --- | --- | --- |
| P<sub>hmtB</sub> | *M. thermautotrophicus* ΔH (MTH_254) | Histone promoter | (Fink et al., 2021) |
| P<sub>synth</sub> | Synthetic variant of P<sub>hmtB</sub> | P<sub>hmtB</sub> variant lacking B-recognition element (BRE) sequence | (Fink et al., 2021) |
| P<sub>synth(BRE)</sub> | Variant of P<sub>synth</sub> | P<sub>synth</sub> variant with B-recognition element (BRE) sequence | (Fink et al., 2021) |
| P<sub>mcrB</sub> | *M. thermautotrophicus* ΔH (MTH_1168) | Methyl-coenzyme M reductase promoter | This study |
| P<sub>ehaA</sub> | *M. thermautotrophicus* ΔH (MTH_384) | Energy-converting hydrogenase complex Eha subunit A promoter | This study |
| P<sub>ehbA</sub> | *M. thermautotrophicus* ΔH (MTH_1251) | Energy-converting hydrogenase complex Ehb subunit A promoter | This study |
| P<sub>slp</sub> | *M. thermautotrophicus* ΔH (MTH_719) | Putative S-layer protein promoter | This study |

**Figure 4.**
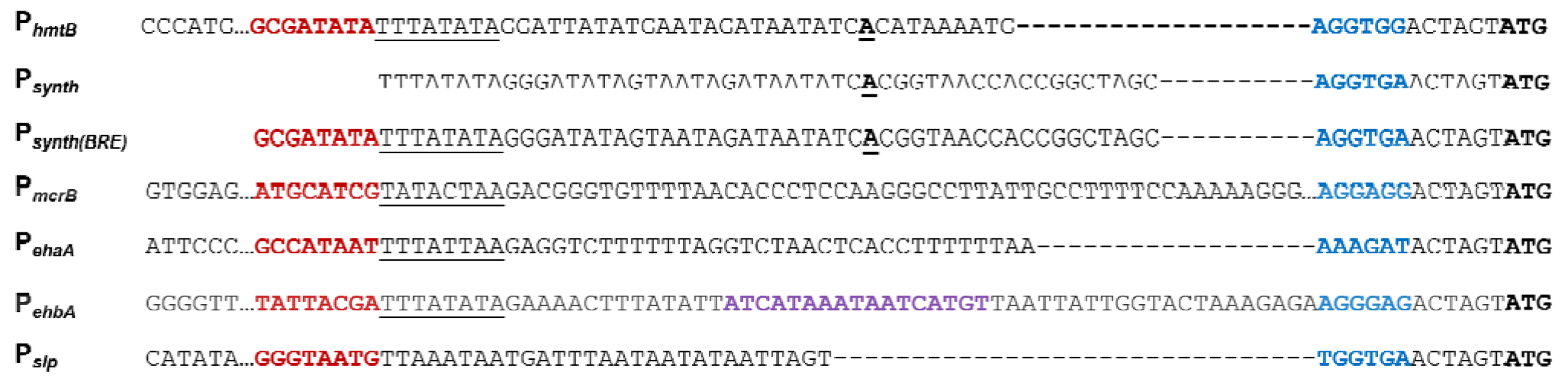
Sequence alignment of promoters tested in FAST assays. **Red/bold**, putative BRE sequences; <u>underlined</u>, putative TATA box sequences; **<u>bold/underlined</u>**, transcriptional start site, confirmed (Darcy et al., 1999; Santangelo et al., 2008); **purple/bold**, putative regulatory element (Kato et al., 2008); **blue/bold**, putative ribosomal binding sites; **bold**, start codon.

**Figure 5.**
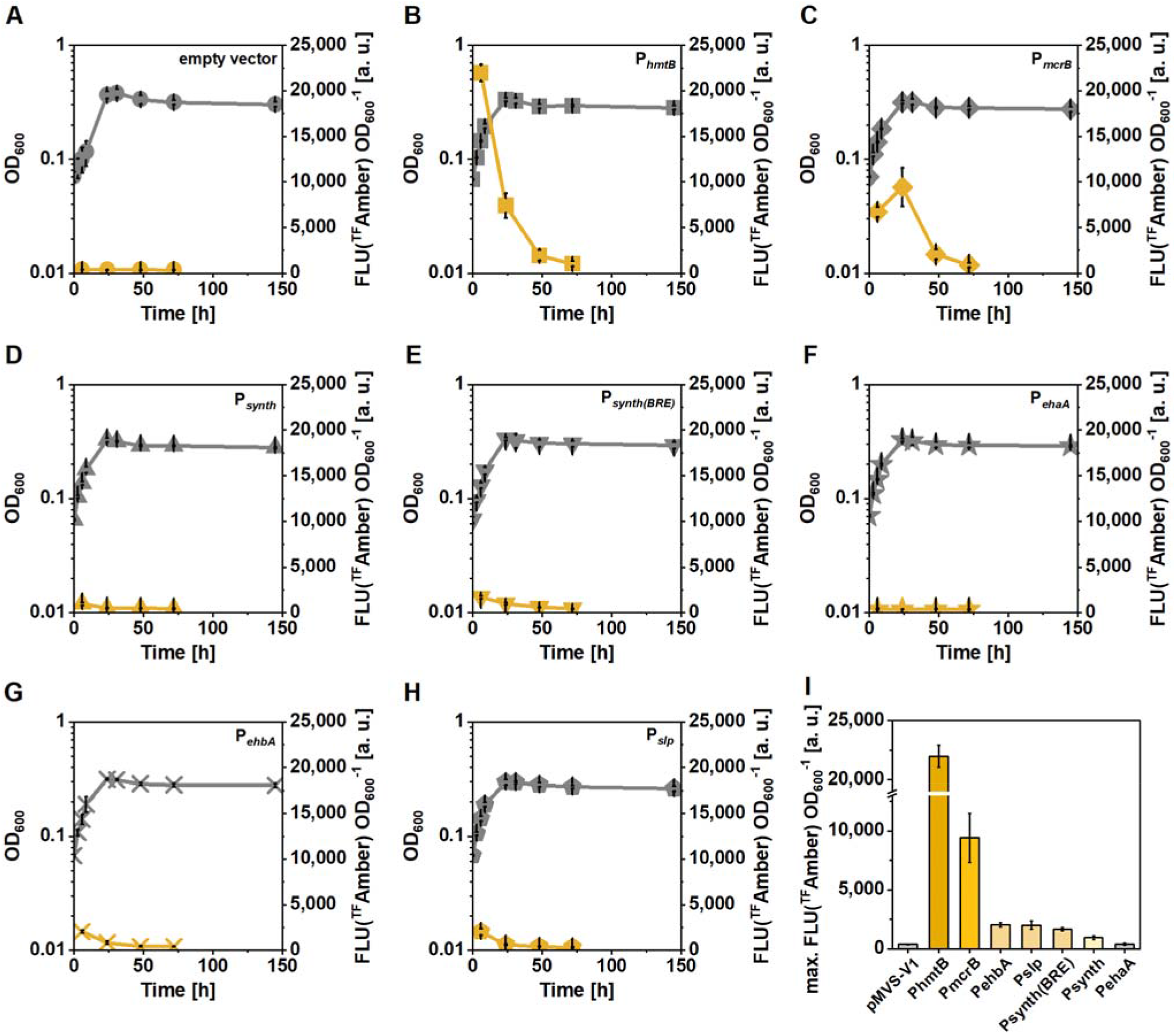
Characterization of different promoters in *M. thermautotrophicus* ΔH using pFAST at 50°C. All strains carried pMVS-V1-based plasmids with the pFAST gene under control of different promoters. **(A)**, pMVS-V1 (empty-vector negative control); **(B)**, P_*hmtB*_; **(C)**, P_*mcrB*_; **(D)**, P_*synth*_; **(E)**, P_*synth(BRE)*_; **(F)**, P_*ehaA*_; **(G)**, P_*ehbA*_; **(H)**, P_*slp*_; **(I)**, maximum fluorescence intensities (^TF^Amber) of pFAST-producing *M. thermautotrophicus* strains. Results from statistical analysis are provided in **Table 4**. Grey, growth (OD_600_); yellow, fluorescence intensity (^TF^Amber) normalized to OD_600_ (FLU(^TF^Amber) OD _600_ ^−1^). Error bars indicate standard deviations, N=3.

Similarly to the previous experiments, the empty-vector negative control did not result in considerable fluorescence (**Figure 5A**). We again confirmed that P_*hmtB*_ resulted in fluorescence only in the early exponential phase, with lower fluorescence intensity during the late exponential, and further reduced fluorescence intensity during the stationary phase (**Figure 5B**). In contrast, P_*mcrB*_ showed lower fluorescence intensity during the early exponential phase, peaked during the late exponential phase, and showed a decline during the stationary phase (**Figure 5C**), which is a distinctly different expression profile compared to P_*hmtB*_. Two synthetic variants of the P_*hmtB*_ promoter, P_*synth*_ (without the B-recognition element [BRE] sequence) and P_*synth(BRE)*_, resulted in a similar expression profile compared to P_*hmtB*_, but with markedly reduced activities (**Figure 5D, E**). Furthermore, we compared the promoters of the two gene clusters encoding the energy-converting hydrogenase isoenzymes, Eha and Ehb (*i*.*e*., P_*ehaA*_ and P_*ehbA*_). While P_*ehbA*_ resulted in a similar expression profile as P_*synth(BRE)*_, P_*ehaA*_ did not seem to be active under the tested conditions (**Figure 5F, G**). The promoter of a putative S-Layer protein, P_*slp*_, was also active with a similar expression profile as P_*synth(BRE)*_ (**Figure 5H**). The maximum fluorescence intensity was significantly higher for P_*hmtB*_ compared to all other promoters, including P_*mcrB*_, which itself had a considerably higher maximum fluorescence intensity compared to the other promoters, but this was not significant in this experiment (**Figure 5I, Table 4**). P_*synth(BRE)*_, P_*ehbA*_, and P_*slp*_ all had similar maximum fluorescence intensities. Those values were significantly higher compared to P_*synth*_ for P_*synth(BRE)*_ and P_*ehbA*_, but not for P_*slp*_ (**Figure 5I, Table 4**). Lastly, P_*ehaA*_ did not reach a fluorescence intensity that was significantly different to the autofluorescence of the cells of the empty-vector negative control (**Figure 5I, Table 4**). To assess the effectiveness of the FAST system further, and as an additional confirmation of the observed expression profiles, we performed the experiment with ^TF^Lime, which led to similar results (**Supplementary Figure 2, Supplementary Table 2**).

**Table 4.** Statistical comparison of different promoters (p-values) for maximum fluorescence intensities shown in Figure 5. Red, highly significant (p ≤ 0.005); orange, significant (p ≤ 0.05); yellow, not significant (p ≤ 0.5); green, not significant (p ≥ 0.5).

|  | pMVS-V1 | P<sub>hmtB</sub> | P<sub>mcrB</sub> | P<sub>ehbA</sub> | P<sub>slp</sub> | P<sub>synth(BRE)</sub> | P<sub>synth</sub> | P<sub>ehaA</sub> |
| --- | --- | --- | --- | --- | --- | --- | --- | --- |
| pMVS-V1 | - | 0.003 | 0.075 | 0.016 | 0.068 | 0.016 | 0.112 | 1.000 |
| P<sub>hmtB</sub> | 0.003 | - | 0.020 | 0.002 | 0.001 | 0.003 | 0.002 | 0.003 |
| P<sub>mcrB</sub> | 0.075 | 0.020 | - | 0.107 | 0.102 | 0.099 | 0.083 | 0.074 |
| P<sub>ehbA</sub> | 0.016 | 0.002 | 0.107 | - | 1.000 | 0.323 | 0.012 | 0.005 |
| P<sub>slp</sub> | 0.068 | 0.001 | 0.102 | 1.000 | - | 0.781 | 0.121 | 0.058 |
| P<sub>synth(BRE)</sub> | 0.016 | 0.003 | 0.099 | 0.323 | 0.781 | - | 0.036 | 0.003 |
| P<sub>synth</sub> | 0.112 | 0.002 | 0.083 | 0.012 | 0.121 | 0.036 | - | 0.076 |
| P<sub>ehaA</sub> | 1.000 | 0.003 | 0.074 | 0.005 | 0.058 | 0.003 | 0.076 | - |

Overall, we demonstrated the functionality of the FAST system with different fluorogens in the thermophilic methanogen *M. thermautotrophicus*. We showed a certain thermolability of the system that will need to be considered for *in-vivo* studies, but which allowed us to investigate the expression profiles of promoters during different growth phases.

## 4 Discussion

We established FAST as a reporter system to enable fluorescence as a read-out in the thermophilic methanogen *M. thermautotrophicus*. We found that FAST is to some extent thermolabile in *M. thermautotrophicus*, and this observation was also previously reported for *T. kivui* (Hocq et al., 2023). We speculate that FAST is either degraded or loses its capacity to result in fluorescence over time due to its thermolability. However, while interpreting our finding that the FAST-mediated fluorescence is not stable over time, it is important to keep in mind that different underlying effects might explain this observation: **1)** a promoter is only active in a certain phase and the observed fluorescence at later time points is solely from still viable FAST protein that is diluted during growth; **2)** FAST is degraded quickly and fluorescence is only measured when a promoter was active during the respective conditions (*e*.*g*., growth phase); **3)** a balance of a certain degree of degradation and newly produced FAST protein defines the actual fluorescence intensity.

For the normalized fluorescence intensity (FLU OD_600_^−1^) this would mean:

- If FAST would be fully thermostable and the promoter would be constitutive, the normalized fluorescence intensity would increase due to accumulation of constantly produced FAST in the cell culture.
- If FAST would be thermostable and the promoter would not be constitutive, the normalized fluorescence intensity would remain stable or slightly decrease because less or no new FAST would be produced and cell division would dilute the amount of available FAST.
- If FAST would be thermolabile and the promoter would not be constitutive, the normalized fluorescence intensity would decrease with a faster rate, because degradation and dilution would happen simultaneously, and less or no new FAST would be produced.

In fact, the second situation is what we had observed with the thermostable β-galactosidase previously (Fink et al., 2021). The P_*hmtB*_ promoter resulted in stable but not further increasing fluorescence over time, which was especially apparent, when normalized to the OD_600_ (*i*.*e*., Miller units; *Figure S7 within* Fink et al., 2021). In the previous work, we had interpreted this finding for P_*hmtB*_ as being a constitutively active promoter because the signal remained stable; however, we did not consider the dilution effects. Here, we observed a marked decrease of the normalized fluorescence intensity, indicating that FAST is degraded and simultaneously diluted during growth, while no or less new FAST is produced (third situation). We interpret our findings in a way that the degradation rate at 60°C was higher than at 50°C. Furthermore, during the late exponential phase the degradation rate of FAST at 60°C was higher than the production rate, while at 50°C the degradation rate was lower than the production rate. Future investigations will have to clarify the actual degradation rate of FAST at the different temperatures. Furthermore, a higher time-resolution in the first 24 hours would help to understand the expression profiles in more detail.

Taken together, our findings here demonstrate that the P_*hmtB*_ promoter is only highly active in the exponential phase and fluorescence intensity drops markedly already in the late exponential phase, which means that P_*hmtB*_ should not be considered constitutive anymore. Further evidence that such regulation patterns are not an artifact of dilution during growth *alone* is provided by the P_*mcrB*_ expression profile. Here, initially lower fluorescence intensity is detected in the early exponential phase, while this increases in the late exponential phase and drops again in the stationary phase. This can only be explained by a sufficiently high protein production rate during the late exponential phase that outweighs the degradation and dilution effects.

For the two synthetic variants of the P_*hmtB*_ promoter, P_*synth*_ and P_*synth(BRE)*_, we confirm our previous finding from the β-galactosidase assays (Fink et al., 2021), for which we found that those promoters result in significantly lower promoter strength and that the presence of the BRE sequence increases the activity slightly. Nevertheless, both promoters showed a (significantly lower but) similar expression profile as P_*hmtB*_, and thus also should not be considered constitutive anymore. Furthermore, we included the promoter of a putative S-layer protein (P_*slp*_) in our study. While there is no report of an S-layer in *M. thermautotrophicus* in the peer-reviewed literature, we find that P_*slp*_ is active at low levels, which warrants further investigation.

Interestingly, we found activity for P_*ehbA*_ but not for P_*ehaA*_ in our conditions. Those promoters were derived from the two isoforms of the energy-converting hydrogenase (Eha and Ehb)-encoding gene clusters (Kaster et al., 2011; Lie et al., 2012). It has been shown that the pH_2_ influences the expression levels of those complexes, and that the Eha-encoding genes were expressed at much lower levels when H_2_ was not limiting (Tersteegen and Hedderich, 1999). A putative regulatory sequence was predicted within the promoter region for P_*ehbA*_ based on microarray data (Kato et al., 2008). This sequence was also found for *ehaA*, but in this case it was located in the beginning of the coding sequence of *ehaA* instead of the promoter region (Kato et al., 2008). From gene deletion studies with the mesophilic methanogen *M. maripaludis*, different physiological roles of the two complexes have been proposed, with Ehb being involved in anabolic functions, while the function of Eha is required for catabolism (Porat et al., 2006; Major et al., 2010; Costa et al., 2013; Lohner et al., 2014). The function of Eha and Ehb in *M. thermautotrophicus* have not been unequivocally deduced.

Different regulatory patterns have been also described for the two isoforms of the methyl-coenzyme M reductase (Mcr and Mrt)-encoding gene clusters (Morgan et al., 1997; Reeve et al., 1997). We did not include P_*mrt*_ here, but we had found that P_*mrt*_ was not active under similar conditions with the β-galactosidase assay before (Fink et al., 2021). In contrast, P_*mcrB*_ resulted in high activity levels with FAST, and showed an expression profile, which would match with higher activities at lower pH_2_. Importantly, instead of typical BRE and TATA-box sequences of archaeal promoters, the P_*mrt*_ promoter contains palindromic sequence repeats, which are rich in thymine and adenine bases, and which overlap with the transcription start site (Shinzato et al., 2008). In addition to the putative regulatory elements discussed above, one such palindromic sequence with high sequence similarity is also present in P_*ehaA*_, which is in general A/T-rich with several non-palindromic repeat sequences. This similarity may indicate similar regulatory mechanisms. Eha and Ehb are membrane-associated proteins, and thus much lower expression levels are reasonable.

Overall, our results support the previous findings of pH_2_-dependent regulation for those promoters, while the responsible mechanisms await further investigation. An important caveat in our experiments is that the pH_2_ drops over time during the batch cultivation in serum bottles. While the lower pH_2_ towards the end of the experiment might trigger certain promoter activities, growth in general ceases and fewer new proteins (including the FAST) are produced. Thus, to investigate effects from pH_2_ on different promoters, the FAST system should be combined with bioreactors, which would allow to set different pH_2_ while keeping the culture in a continuously growing state (Zipperle et al., 2026). Here, we provide a valuable tool that amends the genetic toolbox for *M. thermautotrophicus* and potentially other thermophilic methanogens, such as *Methanothermobacter marburgensis*, for which genetic tools have recently been reported (Klein et al., 2025; Unger et al., 2026). Future use in cell biology is possible but an improvement of thermostability would be beneficial and should be aimed for by applying targeted mutagenesis or adaptive laboratory evolution with FAST, as demonstrated for *T. kivui* (Hocq et al., 2023).

## 7 Author Contributions

Tina Baur (T.B.) and Bastian Molitor (B.M.) designed the experiments. T.B. performed all experimental work. T.B. and B.M. analyzed the data. T.B. and B.M. prepared figures and tables. Statistical analyses were done by B.M. and Maximilian Flaiz (M.F.). B.M. and L.T.A. supervised the work and acquired funding. M.F. drafted the introduction, T.B. drafted the Materials and Methods, and B.M. drafted the remaining manuscript. All authors contributed to editing and approved the final version.

## 9 Acknowledgments

n/a

## 10 Supplementary Material

Supplementary Materials were uploaded separately on submission.

## 11 Data Availability Statement

The datasets for this study can be provided by the corresponding author.

