## Supplementary Material for "Establishing the fluorescence-activating and absorption-shifting tag as a fluorescent reporter protein in *Methanothermobacter thermautotrophicus* ΔH"

### 1.1 Supplementary Figures

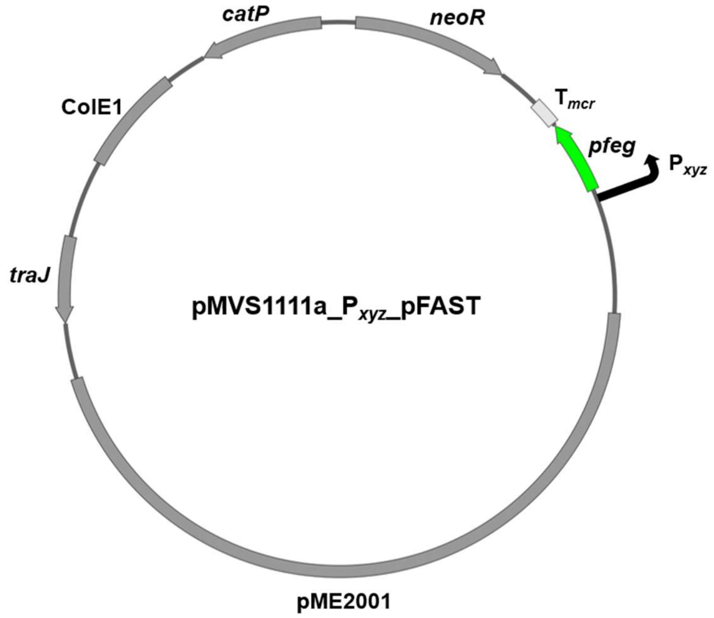


**Supplementary Figure 1. Plasmid map of pFAST-encoding shuttle vector for *M. thermautotrophicus* ΔH.** pME2001, replicon for *M. thermautotrophicus*; *traJ*, gene encoding conjugal transfer function; ColE1, Gram-negative replicon for *E. coli*; *catP*, chloramphenicol resistance gene; *neoR*, neomycin resistance gene; $P_{xyz}$, promoter to control pFAST expression ($P_{xyz}$= $P_{hmtB}$, $P_{synth}$, $P_{synth(BRE)}$, $P_{mcrB}$, $P_{ehaA}$, $P_{ehbA}$, $P_{slp}$); *pfeg*, pFAST-encoding gene codon-optimized for *M. thermautotrophicus*; $T_{mcr}$, terminator sequence of *mcr* operon of *Methanococcus voltae*.

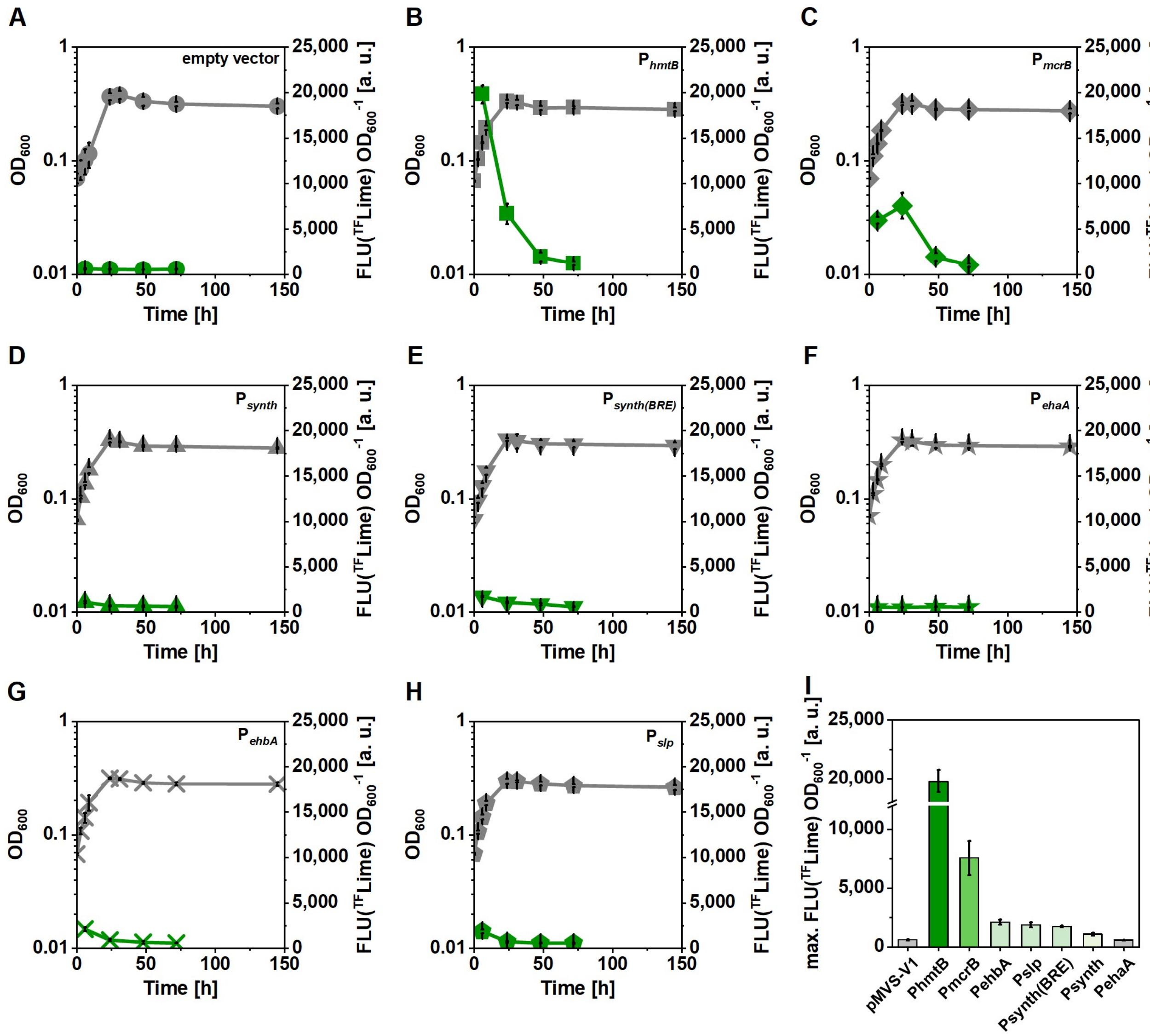


**Supplementary Figure 2. Characterization of different promoters in *M. thermautotrophicus* ΔH using pFAST at 50°C.** All strains carried pMVS-V1-based plasmids with the pFAST gene under control of different promoters. **(A)**, pMVS-V1 (empty-vector negative control); **(B)**, $P_{hmtB}$; **(C)**, $P_{mcrB}$; **(D)**, $P_{synth}$; **(E)**, $P_{synth(BRE)}$; **(F)**, $P_{ehaA}$; **(G)**, $P_{ehbA}$; **(H)**, $P_{slp}$; **(I)**, maximum fluorescence intensities ($^{TF}$Lime) of pFAST-producing *M. thermautotrophicus* strains. Grey, growth ($OD_{600}$); green, fluorescence intensity ($^{TF}$Lime) normalized to $OD_{600}$ (FLU($^{TF}$Lime) $OD_{600}^{-1}$). Error bars indicate standard deviations, N=3.

### 1.2 Supplementary Tables

**Supplementary Table 1. Full sequences of codon-optimized pFAST and tdpFAST genes, and all promoters characterized in this study using pFAST.** Red sequence for tdpFAST shows the GGGSGGG-linker-encoding sequence.

| Promoter | Sequence |
|---|---|
| pFAST | ATGGAGCACGTCGCTTTTGGATCTGAGGATATTGAGAACACACTTGCCAA<br>TATGGATGATGAACAGCTTGATAGGCTAGCCTTCGGGGTAATTCAGCTGG<br>ACGGTGATGGAAACATCTTATTGTATAATGCCGCGGAAGGGGACATAACG<br>GGGCGAGACCCTAAACAAGTTATCGGTAAGAACTTTTTCAAAGACGTAGC<br>TCCTGGGACCGATACCCCCGAGTTCTATGGAAAATTTAAAGAAGGCGCAG<br>CATCCGGAAACCTGAATACCATGTTTGAATGGACAATACCAACTTCAAGA<br>GGTCCGACTAAAGTGAAGGTTCATCTCAAGAAGGCACTCAGTGGCGATAG<br>GTACTGGGTCTTCGTGAAGCGGGTGTGA |
| tdpFAST | ATGGAACATGTAGCCTTCGGCAGTGAGGACATTGAAAACACTTTAGCCAA<br>CATGGACGACGAACAGCTTGATAGATTAGCTTTCGGAGTTATACAGTTAG<br>ATGGTGACGGGAACATTCTCCTGTATAATGCCGCCGAGGGAGACATCACA<br>GGTAGGGACCCTAAGCAAGTCATAGGAAAGAACTTCTTTAAAGATGTAGC<br>GCCAGGAACTGATACCCCGGAATTCTACGGTAAGTTCAAAGAGGGTGCCG<br>CGTCAGGTAACCTGAACACGATGTTCGAGTGGACAATTCCGACTTCAAGG<br>GGACCCACGAAAGTGAAAGTACACCTCAAAAAGGCACTTTCAGGAGATC<br>GGTACTGGGTCTTTGTCAAGAGAGTG**GGTGGAGGGTCCGGTGGTGGC**G<br>AACATGTCGCTTTCGGATCTGAGGATATCGAGAACACACTTGCCAATATG<br>GATGACGAACAACTGGATAGACTCGCATTCGGGGTAATACAGCTGGATGG<br>GGACGGAAATATCCTGCTCTATAATGCTGCAGAAGGCGACATAACAGGCC<br>GGGATCCCAAGCAGGTTATCGGGAAAAATTTCTTTAAAGATGTTGCACCA<br>GGGACAGATACCCCTGAGTTTTATGGGAAATTTAAGGAAGGGGCTGCATC<br>TGGCAATTTGAATACCATGTTTGAGTGGACCATCCCCACCTCCAGGGGGC<br>CTACCAAGGTGAAAGTTCACTTGAAGAAGGCATTGAGCGGCGATAGGTAT<br>TGGGTGTTTGTTAAAAGGGTGTGA |
| $P_{hmtB}$ | CCCATGAACCAACCGATGGCTCAGAAAAACCTTAAAATTAGCGATATATT<br>TATATAGGATTATATGAATAGATAATATCACATAAAATGAGGTGG |
| $P_{synth}$ | TTTATATAGGGATATAGTAATAGATAATATCACGGTAACCACCGGCTAGC<br>AGGTGA |
| $P_{synth(BRE)}$ | AGCGCGATATATTTATATAGGGATATAGTAATAGATAATATCACGGTAAC<br>CACCGGCTAGCAGGTGA |
| $P_{mcrB}$ | GTGGAGAGCTCTAAATATAATTTATTACCGGCTCTGGACCCATGGGGCAG<br>TGACTGTGAATGACCAGAGGATGATTATTATTACTCCCTGCTGATGATACT<br>CTGCTTTGGGATGATCACAAAAATCAACGCCACCATATAAAAAATCCAGA<br>AAACCCAAAAAAAGGTGTTTTTACATATAAAGCTTTCGACTTGATTCAGA<br>AAGAAATCTTAATTAATTATAATCAAGTTAAATATGCATCGTATACTAAG<br>ACGGGTGTTTTAACACCCTCCAAGGGCCTTATTGCCTTTTCCAAAAAGGG<br>GGGTCCTGGATGTTTAGTATACGGAAATGATTGTTAGGAGG |

| | |
|---|---|
| $P_{ehaA}$ | ATTCCCACCCCATCATGTCCTGGAGGTTCCTATGGAGTCACATCCACTTAC CGGTTTCTGTGAATATAATCACGAAAACGACTATGAAAATCCTTAAAATT TATTACTAAATCTTCTGTAGAGGGACCCTTCAGATATCAGTAAAAAAAAG TTTCTGTAGAGAAACCTCCAGATATTAGTAAAAAAGGCCATAATTTTATT AAGAGGTCTTTTTTAGGTCTAACTCACCTTTTTTAAAAAGAT |
| $P_{ehbA}$ | GGGGTTATTACGGCAGCCTCTTCCCCTTCAAGTGTTTAATCATCGGATCCC CTATTTTATTACGATTTATATAGAAAACTTTATATTATCATAAATAATCAT GTTAATTATTGGTACTAAAGAGAAGGGAGC |
| $P_{slp}$ | CATATAAGAATTTCATCCAGGGGTTATATCTTTTTTATGAAGCAGTTCTTC TAATTGTCCCACTTCAAATCTTTTTATATGTTTTTTTTATTCGCCGTTAAGA TGATGCTATTTTTTTTGGTTTAGGGTAATGTTAAATAATGATTTAATAATA TAATTAGTTGGTGA |

**Supplementary Table 2. Maximum fluorescence values (FLU $OD_{600}^{-1}$ [a.u.]) for all experiments.**

| | | Fluorogen | | |
|---|---|---|---|---|
| **FAST variant** | **promoter** | **$^{TF}$Lime** | **$^{TF}$Amber** | **$^{TF}$Coral** |
| ***Temperature comparison*** | | | | |
| ***50°C*** | pMVS-V1 | 465.0 ± 15.3 | | |
| pFAST | $P_{hmtB}$ | 16592.2 ± 793.4 | | |
| tdpFAST | $P_{hmtB}$ | 13991.4 ± 583.4 | | |
| ***60°C*** | pMVS-V1 | 513.7 ± 48.8 | | |
| pFAST | $P_{hmtB}$ | 8063.2 ± 274.7 | | |
| tdpFAST | $P_{hmtB}$ | 4362.2 ± 414.6 | | |
| ***Fluorogen comparison*** | | | | |
| | pMVS-V1 | 495.9 ± 20.5 | 331.6 ± 50.4 | 414.3 ± 53.5 |
| pFAST | $P_{hmtB}$ | 10320.5 ± 1427.0 | 11316.4 ± 1174.0 | 5970.7 ± 768.7 |
| ***Promoter comparison*** | | | | |
| | pMVS-V1 | 601.3 ± 60.9 | 395.6 ± 17.4 | |
| pFAST | $P_{hmtB}$ | 19786.3 ± 969.5 | 21977.8 ± 935.2 | |
| pFAST | $P_{mcrB}$ | 7567.0 ± 1434.0 | 9417.2 ± 2089.2 | |
| pFAST | $P_{ehbA}$ | 2109.9 ± 205.2 | 2042.4 ± 177.5 | |
| pFAST | $P_{slp}$ | 1875.6 ± 223.4 | 2020.5 ± 361.0 | |
| pFAST | $P_{synth(BRE)}$ | 1730.2 ± 64.0 | 1684.9 ± 141.2 | |
| pFAST | $P_{synth}$ | 1091.3 ± 113.0 | 960.5 ± 164.5 | |
| pFAST | $P_{ehaA}$ | 586.0 ± 28.2 | 392.2 ± 88.4 | |
